# Salicin modifies osteoarthritis progression by binding on IRE1α and inhibiting IRE1α mediated endoplasmic reticulum stress

**DOI:** 10.1101/2022.04.06.487311

**Authors:** Zhenglin Zhu, Shengqiang Gao, Cheng Chen, Wei Xu, Pengcheng Xiao, Zhiyu Chen, Chengcheng Du, Bowen Chen, Yan Gao, Chunli Wang, Junyi Liao, Wei Huang

## Abstract

**Objectives:** To investigate the effect and mechanisms of salicin (SA) on osteoarthritis (OA) progression.

**Methods:** Primary rat chondrocytes were stimulated with TNF-α and treated with or without SA. CCK-8 was utilized to determine the cytotoxicity of SA. RT-qPCR, Western Blotting and immunofluorescence staining were used to detect inflammatory factors, cartilage matrix degeneration markers, cell proliferation and apoptosis markers expression at mRNA and protein levels respectively. EdU assay and flow cytometer analysis were utilized for evaluating cell proliferation and apoptosis. RNA-sequencing, molecular docking, drug affinity responsive target stability and WB were applied to clarify mechanisms. Rat OA model was used to evaluate the effect of intra-articular injection of SA on OA progression.

**Results:** No obvious cytotoxicity was found with the treatment of 10 μM SA. SA rescued TNF-α induced degeneration of cartilage matrix, inhibition of chondrocytes proliferation, and promotion of chondrocytes apoptosis. In mechanism, we clarified SA could directly bind on IRE1α and occupy IRE1α phosphorylation site, followed with inhibiting IRE1α phosphorylation and regulating IRE1α mediated endoplasmic reticulum (ER) stress by IRE1α-IκBα-p65 signaling. Finally, intra-articular injection of SA loaded PLGA could ameliorate OA progression by inhibiting IRE1α mediated ER stress in OA model.

**Conclusions:** SA alleviates OA by directly binding on ER stress regulator IRE1α and inhibits IRE1α mediated ER stress by IRE1α-IκBα-p65 signaling. Topical use of small molecular drug SA holds the potential of modifying OA progression.

## 1. Introduction

Osteoarthritis (OA) is the most prevalent joint disease which affects an estimated more than 500 million people worldwide^1^. Exploring disease-modifying OA drugs (DMOADs) for modifying the structural progression of OA is considered a potential strategy for the treatment of early or middle stage OA^2,3^.

Willow bark is regarded as one of the successful examples of the modern drug developed from herbal remedy, which is originally recognized around two hundred years ago. Salicin (SA) is the main chemically standardized willow bark extract, its chemical oxidation resulted in a new substance termed “salicylic acid”, and the acetylated derivative is finally turned into the famous drug called “Aspirin”^4,5^. SA is metabolized into salicylic acid after oral administration, and then plays roles in the treatment of pain, headache, and inflammatory conditions^5,6^. However, the formation of salicylic acid alone is unlikely to explain analgesic or anti-rheumatic effects of willow bark^7^, which indicates potential mechanisms that need to be further clarified. On the other hand, as a small molecule drug, SA is characterized by easy to manufacture, absorb and can cross cell membrane directly. Recently, studies reported SA held anti-inflammatory effects and further presented inhibiting angiogenesis effects^8^, prevented cellular senescence^9^, and exhibited anti-irritation and anti-aging effects in dermatological applications^10^. However, based on our knowledge, the effects and mechanisms of SA on OA cartilage degeneration have not been studied.

In the present study, we investigated the effects of SA on cartilage degeneration by both in *vitro* and in *vivo* tests. We clarified that SA prevented cartilage degeneration by alleviating the function of inositol-requiring enzyme 1α (IRE1α) signaling mediated endoplasmic reticulum (ER) stress. Our findings indicated the potential therapeutic effects of SA on OA progression by intra-articular injection.

## 2. Materials and Methods

### 2.1 Chondrocyte culture and chemicals

Experiment protocols were approved by the Institutional Review Board (IRB) of Chongqing Medical University (NO.2020-018). Male Sprague-Dawley (SD) rats (n=30) were fed in the Specific Pathogen Free animal facilities. Primary rat knee chondrocytes were isolated from articular cartilage of 4-day-old neonatal rat, as described previously^11^. Each treatment group has three rats, and the chondrocytes from each rat was subjected to following experiments. Passage 0 or 1 chondrocytes were subjected to following experiments.

SA (Selleck Chemicalsxz, S2351, TX, USA) was dissolved in 0.1% dimethylsulfoxide (DMSO), and 0.1% DMSO was used as control. Tumor necrosis factor α (TNF-α) was purchased from Peprotech (400-14, NJ, USA), 4-Phenylbutyric acid (4-PBA) was purchased from Med Chem Express (Shanghai, China). Unless indicated otherwise, all chemicals were purchased from Sigma-Aldrich or Corning.

### 2.2 Cell viability

Cell viability was determined by Cell Counting Kit-8 (CCK-8, Med Chem Express, Shanghai, China) following the manufacturer’s protocol. For the toxicity of dosage, chondrocytes were treated with SA in gradient concentration (0-100 μM) for 48 hours. For the sustained toxicity of SA, chondrocytes were cultured in a medium containing SA in different concentrations for 5 days. The optical density (OD) values were determined at a wavelength of 450 nm. The non-linear regression analysis was used to calculate the half-maximal inhibitory concentration (IC50) values (percent cell proliferation versus concentration).

### 2.3 Reverse transcription-quantitative polymerase chain reaction (RT-qPCR)

Total RNA was extracted from the chondrocytes with TRIZOL reagent (Thermo Fisher Scientific, MA, USA) according to the manufacturer’s instructions. Reverse transcriptions were carried out by using EvoScript Universal cDNA Master Reagent Kit (Med Chem Express, Shanghai, China). The SYBR Green qPCR Master Mix (Med Chem Express, Shanghai, China), cDNA and primer were mixed according to the manufacturer’s instructions. QPCR reaction protocols were as follows: 5 minutes at 95°C, 40 cycles of 10 seconds at 95°C, 20 seconds at optimal temperature for each pair of the primers, and 20 seconds at 72°C respectively. Melting curves were generated at every endpoint of amplification for 10 seconds at 95°C before 30 increments of 0.5°C from 65 to 95 °C. GAPDH was used as a reference gene. All sample values were normalized to GAPDH expression by using 2^−ΔΔCT^ method. All qPCR were performed with three independent experiments. The qPCR primer sequences are shown in Table 1. Table 1. List of RT-qPCR primers.

**Table 1.**
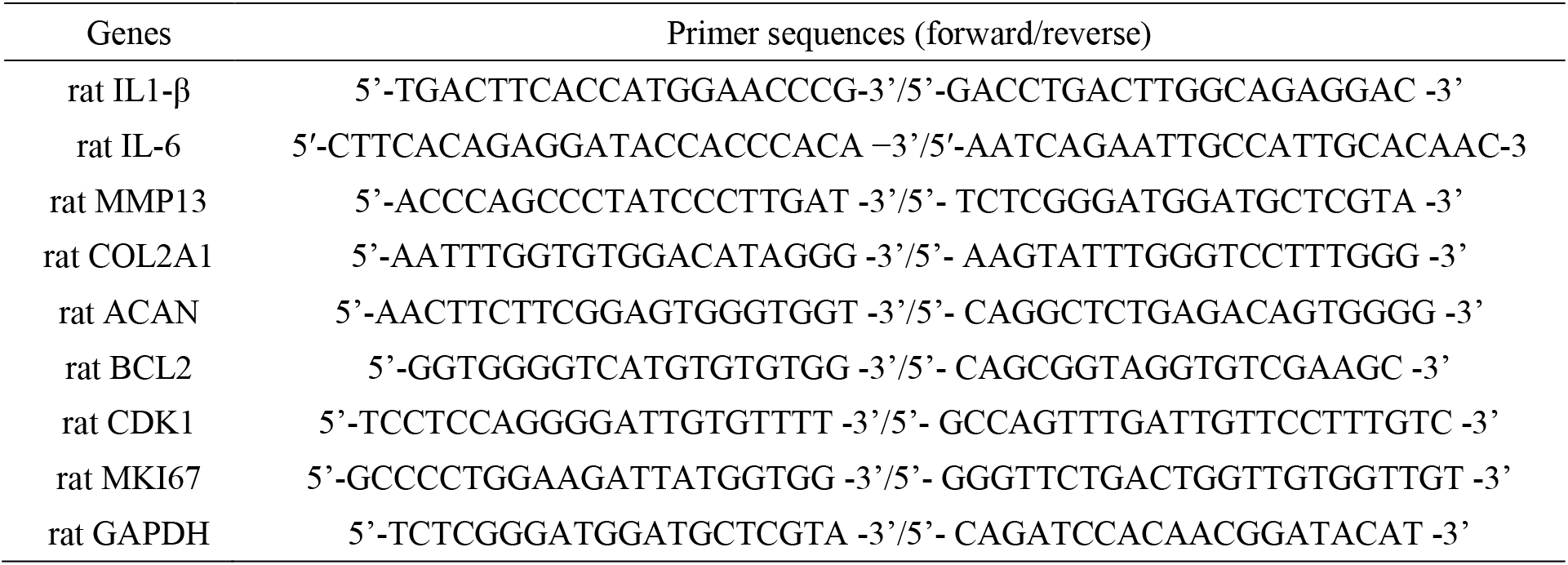
List of RT-qPCR primers.

### 2.4 Western blotting (WB)

Chondrocytes treated with or without SA were lysed in cold Radioimmunoprecipitation (RIPA, Beyotime, Shanghai, China) buffer containing phosphatase and protease inhibitor, then centrifuged at 13000 g at 4 °C for 15minutes. Equivalent quantity of protein (30 μg) in each group was separated by 10-12% SDS-polyacrylamide gel electrophoresis (SDS-PAGE) and transferred to a polyvinylidene difluoride (PVDF) membrane (Roche, Basel, Switzerland). As for detection aggrecan, samples were deglycosylated with manufacturer’s instructions. After blocking with 5% skimmed milk, membranes were probed with primary antibodies against Col2α1 (Abcam, ab34712), Aggrecan (Abcam, ab3773), MMP13 (Zen Bio, 820098), BCL2 (Zen Bio, R22494), CDK1 (Abcam, ab134175), total-ERK1/2 (CST, #4695), phospho-ERK1/2 (CST, #4370), total-IκBα (CST, #4812), phospho-IκBα (CST, #2859), phospho-IRE1α (Zen Bio, R26310), IRE1α (bs-16696R), GRP78 (Zen bio, R24509), NF-κB p65 antibody (CST,#8242), Phospho-NF-κB p65 antibody (CST, #3033), GAPDH (Zen Bio, 380626) overnight at 4 °C. Horseradish peroxidase-conjugated anti-mouse or anti-rabbit IgG (Abcam, ab6728, ab6721) was used as secondary antibodies. Enhanced Chemiluminescence (ECL, Thermo Fisher Scientific, MA, USA) detection system was used to detect the protein bands on the membrane. The intensity of the protein bands was analyzed by Image J software, using GAPDH as reference protein.

### 2.5 Cell immunofluorescence (IF) assays

Cells on coverslips were incubated with 6% normal goat serum and 4% bovine serum albumin in phosphate buffer solution (PBS) for 1 hour at 37°C. Then, cells were incubated with the NF-κB p65 antibody (CST, #8242), Phospho-NF-κB p65 antibody (CST, #3033) overnight at 4°C. Next day, the fluorescein isothiocyanate (FITC/CY3) labeled goat anti-rabbit IgG was used as a secondary antibody and incubated for 1 hour at room temperature. ER-Tracker Red (C1041 Beyotime) was applied according to the manufacture’s protocols. Cells were washed with 1 × PBS for 5 times to clear the dyes. Finally, the cells were counterstained with 4’,6-diamidino-2-phenylindole (DAPI) and observed by fluorescence microscopy.

### 2.6 Cell proliferation assay

Chondrocyte proliferation was quantified by an ethynyl deoxyuridine (EdU) DNA in vitro proliferation detection kit (RiboBio, Guangzhou, China) according to the manufacturer’s instructions. The number of EdU-labelled cells was calculated from fields randomly selected in each well by two independent laboratory technicians in a blinded, random fashion.

### 2.7 Cell apoptosis assay

Chondrocytes in each treatment group were trypsinized, collected and washed three times with cold PBS. About 4 × 10^5^ cells/ml suspensions per group were mixed with the Annexin V-FITC and PI binding buffer for 20 minutes according to the standard protocol of the Annexin V-FITC kit (Beyotime, Beijing, China). Finally, the mixtures were subjected to a flow cytometer (BD Biosciences, CA, USA) for apoptosis analysis.

### 2.8 ELISA

The levels of MMP13 in chondrocyte supernatants were measured using the ELISA kits (E-EL-R0045c, Elabscience, Wuhan, China) according to the manufacturer’s instructions, OD value was recorded at a wavelength of 450 nm.

### 2.9 RNA-sequencing (RNA-Seq) and bioinformatics analysis

Total RNA extracted from chondrocytes treated with or without SA were subjected to RNA-Seq. RNA samples were sent to OE Biomedical Technology Co., Ltd. (Shanghai, China) for RNA-Seq. RNA sequencing was done on the HiSeq2500 system (Illumina, CA, USA). The R language program was used to analyze the raw data and identify differentially expressed genes (DGEs). Selecting differential transcripts with p-values ≤ 0.05 and fold change ≥ 2 were followed with gene ontology (GO) biological function enrichment analysis and Kyoto gene and genome encyclopedia (KEGG) signal pathway enrichment analysis.

### 2.10 Target prediction and molecular docking

SA structure was downloaded from the Pubchem database and converted to mol2 format. The three-dimensional structure of the IRE1α (PDB ID: 6HX1) protein was downloaded from the Research Collaboratory for Structural Bioinformatics (RCSB) protein database. Autodock vina 1.1.2 was used for semiflexible docking with SA and IRE1α. The parameter exhaustiveness was set as 20 to increase calculation accuracy. The best affinity (−8.567 kcal/mol) conformation was selected as the final docking conformation.

### 2.11 Drug affinity responsive target stability (DARTS)

DARTS was done as previously described^12,13^. Briefly, chondrocytes were treated with cold M-PER lysis buffer (Thermo Fisher, MA, USA) containing a protease inhibitor cocktail (1 mM Na3VO4 and 1 mM NaF). Protein lysates were firstly mixed with 10x TNC buffer (500 mM Tris–HCl pH=8.0, 500 mM NaCl and 100 mM CaCl2 at a ratio of 1:1), then subjected to incubate with DMSO or SA for 1 hour at room temperature. Next, sample was proteolyzed in gradient concentrations of pronase (Roche, Basel, Switzerland) for 10 minutes following with adding 2 μl of cold 20x protease inhibitor cocktail. Coomassie blue staining (Beyotime, Shanghai, China) was carried out for estimation of relative abundance of proteins. An equal portion of each sample was loaded onto SDS-PAGE gel for Western blotting.

### 2.12 Animal model

SD rats were randomizely devided into four groups. After anesthesia and standard aseptic surgical procedures ACLT was performed in SD rats’ (200 ± 20 g) right knee to establish an OA model (duration: four weeks) ^14,15^, rats only open knee joints were regarded as a sham group (n=5). Poly (lactic-co-glycolic acid) (PLGA) (Sigma-Aldrich, MO, USA) was dissolved in CH_2_Cl_2_ and mixed with ultrasonic dissolved SA, then PLGA loaded SA was obtained by double-emulsion method as reported previously^16,17^. The animals that received ACLT were randomly divided into three groups: ACLT (PBS only, n=5), vehicle (PLGA vehicle, n=5), and SA (PLGA vehicle with SA, n=5). Four weeks after surgery, intra-articular injection of 100 μL PLGA + SA suspension (the equivalent of 1 mM SA, PLGA loaded), PLGA only and PBS only were done respectively. Four weeks post intra-articular injection, rats were sacrificed and knee joints were harvested.

### 2.13 Histological analysis

Samples were fixed in 4% paraformaldehyde (Beyotime, Beijing, China), decalcified in 0.5 M ethylenediaminetetraacetic acid (EDTA) and embedded in paraffin. Serial 5-μm-thick sections were obtained and subjected to histological and other specialty staining evaluations. Hematoxylin-eosin (H&E) and safranin-O/green stainings were carried out as previously described^18-20^. The articular surface of the femur and tibia in coronal sections of the knee joint was qualitatively and semi-quantitatively analyzed as previously characterized^14,15^. Each specimen was scored by three orthopaedic surgeons following the double-blind principle according to Osteoarthritis Research Society International (OARSI) criterion for rat^21^.

As for Immunohistochemistry (IHC) staining, dewaxed slices were immersed in sodium citrate buffer and heated in gradient temperature for antigen retrieving, soaked in 3% H_2_O_2_ to remove the endogenous peroxidase activity and blocked with caprine serum. Then slices were incubated with primary MMP13 (Zen Bio, 820098), GRP78 (Zen Bio, R24509), p-IRE1α (Zen Bio, 530878) antibody respectively at 4°C overnight. Biotin-labeled secondary antibody (ZSGB-BIO, ZLI-9017) and 3,3′-Diaminobenzidine (DAB) were used for IHC.

As for IF staining, frozen sections were fixed with 4% paraformaldehyde, dehydrated in 30% sucrose solution, permeabilized with 0.1 % Triton X-100 and blocked with 5 % BSA. Then sections were incubated overnight at 4 °C with primary Phospho-NF-κB p65 (CST, #3033) antibody. Sections were dealt with appropriate fluorescein-conjugated secondary antibodies. DAPI was utilized for nuclear staining.

### 2.14 Sample size determinations

For in vitro experiments analyzed as fold changes or quantitative analysis, we calculated a minimum sample size (n ≥3) with three independent experiments. For *in vivo* experiments, based on the previous experiments, we yielding a sample size n ≥ 4 at alpha level of 0.05 and power of 80%^22,23^.

### 2.15 Statistical analysis

All data sets were compared using GraphPad Prism (GraphPad Software, La Jolla, CA) version 9.0. Data are shown as mean ± standard deviation (SD). For in *vivo* and vitro studies, Unpaired Student’s t test (for two groups), one-way ANOVA (for multiple groups) were used followed by the Tukey-Kramer test. Values of *P* < 0.05 were considered statistically significant.

## 3. Results

### 3.1 Phenotypic characterization of primary articular chondrocytes and cytotoxicity of SA on chondrocytes

Primary articular chondrocytes were characterized by Toluidine Blue staining and IF staining with Col2a1(red) and ACAN (green) (Figure 1A). We firstly identified the effects of SA on cell cycle and found that SA promoted chondrocytes proliferation (Figure 1Ba), quantitative analysis (Figure 1Bb) showed SA increased S-phase chondrocytes significantly. Next, cytotoxicity of SA to chondrocytes was assessed. As shown in Figure1C, 5 μM, 10 μM and 20 μM SA statistically promoted chondrocytes proliferation and 10 μM was the highest, however, 50 μM and 100 μM SA dramatically reduced cell viability. Furthermore, we found 10 μM SA treatment promoted chondrocytes proliferation from day 1 to day 4 compared with DMSO group. From these experiments, IC50 of SA was 42.30 μM (Figure 1E). Taken together, 10 μM SA was considered not toxic to chondrocytes and selected as the optimal dose for next investigations.

**Figure 1:**
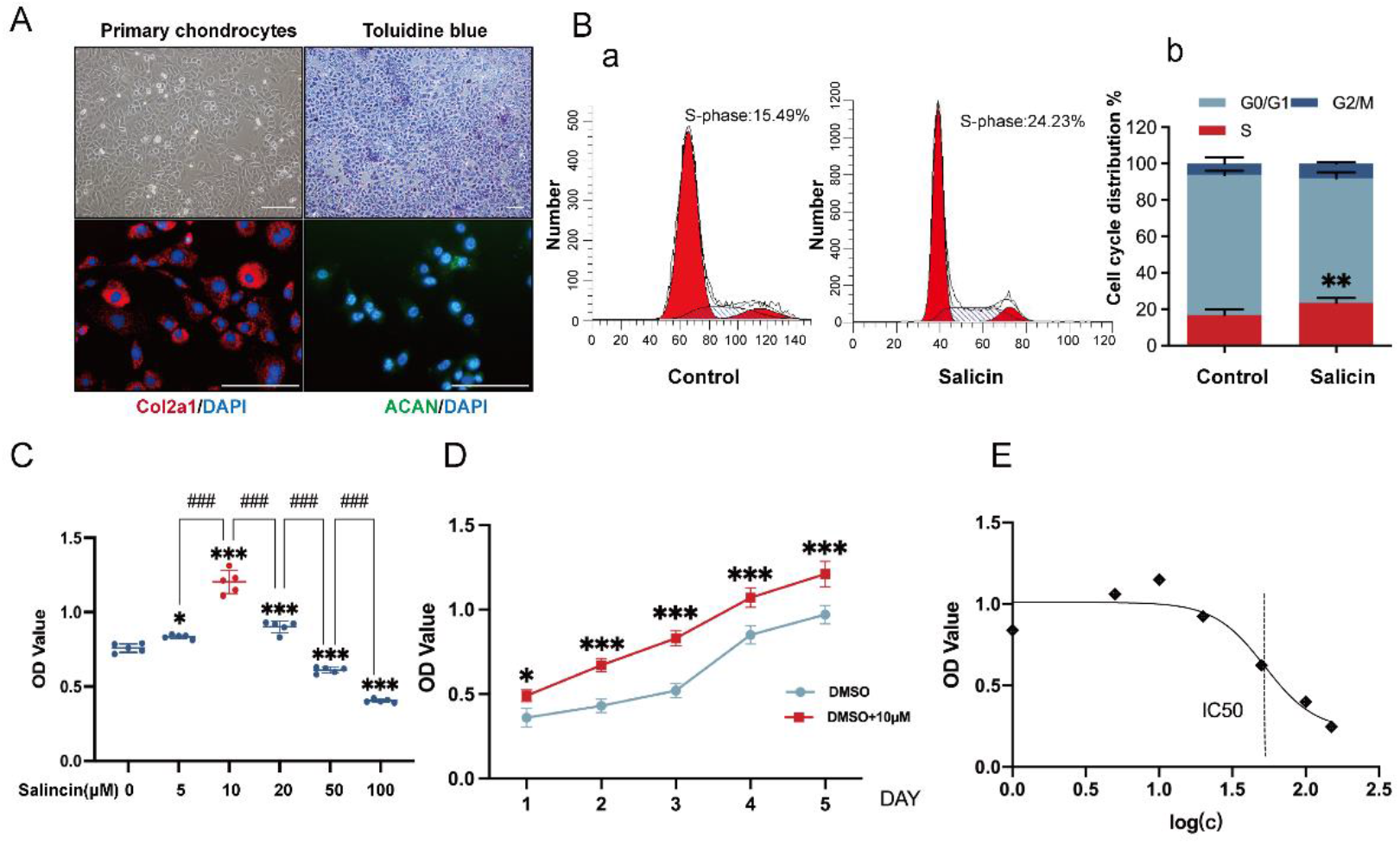
Phenotypic characterization of primary articular chondrocytes and cytotoxicity of SA on chondrocytes. (A) Staining of articular chondrocytes in primary cultures with Toluidine Blue. Immunostaining of primary articular chondrocytes with Col2a1(red), ACAN (green) and DAPI (blue). (B)Flow cytometry for cell cycle progression of chondrocytes with or without SA(a), quantitative analysis(b) found SA promotes S-phase cells significantly (n=3, Student’s t test). (C) Chondrocytes were treated with SA in gradient concentration (0-100 μM) for 48 hrs and subjected to CCK-8 analysis. From 5 μM to 20 μM SA treatment, OD values were statistically higher than 0 μM (DMSO) group, and for 50 μM and 100 μM SA treatment, OD values were statistically lower than 0 μM (DMSO) group (n=5, one-way ANOVA). (D) Chondrocyte proliferation curve of SA treatment from day 1 to day 5, 10 μM SA treatment statistically promoted cell proliferation from day 1 to day 5 compared with DMSO group.(n=5, Student’s t test). (E) IC50 of SA on chondrocytes, concentrations were transferred to Log(c). The data are expressed as mean ±SD, *p<0.05, **p< 0.01, ***p<0.001, * compared with DMSO group, #p<0.05, ##p<0.01, ###p< 0.001, # compared with 10μM group and ns, not significant.

### 3.2 SA inhibits TNF-α induced chondrocytes inflammatory factor expression and extracellular matrix degeneration

We firstly explored the effects of SA on TNF-α induced chondrocytes inflammatory factors and matrix gene expressions. As shown in Figure 2A, TNF-α dramatically increased interleukin-1β (IL-1β), interleukin-6 (IL-6), matrix metalloproteinase 13 (MMP13) gene expression and inhibited Col2a1 and ACAN gene expression in chondrocytes when compared with the control group. However, these effects could be diminished by 48 hours’ treatment of 10 μM SA in *vitro*. What was more, we found TNF-α induced chondrocytes matrix degeneration by elevating the expression and secretion of MMP13 would be reverted by SA at protein level (Figure 2B-C). Meanwhile, we found SA could rescue TNF-α induced cartilage matrix degradation by detecting expressions of Col2a1 and ACAN at protein level (Figure 2C). Quantitative analysis showed the same trend. These results indicated that SA inhibited TNF-α induced chondrocytes degeneration in *vitro*.

**Figure 2:**
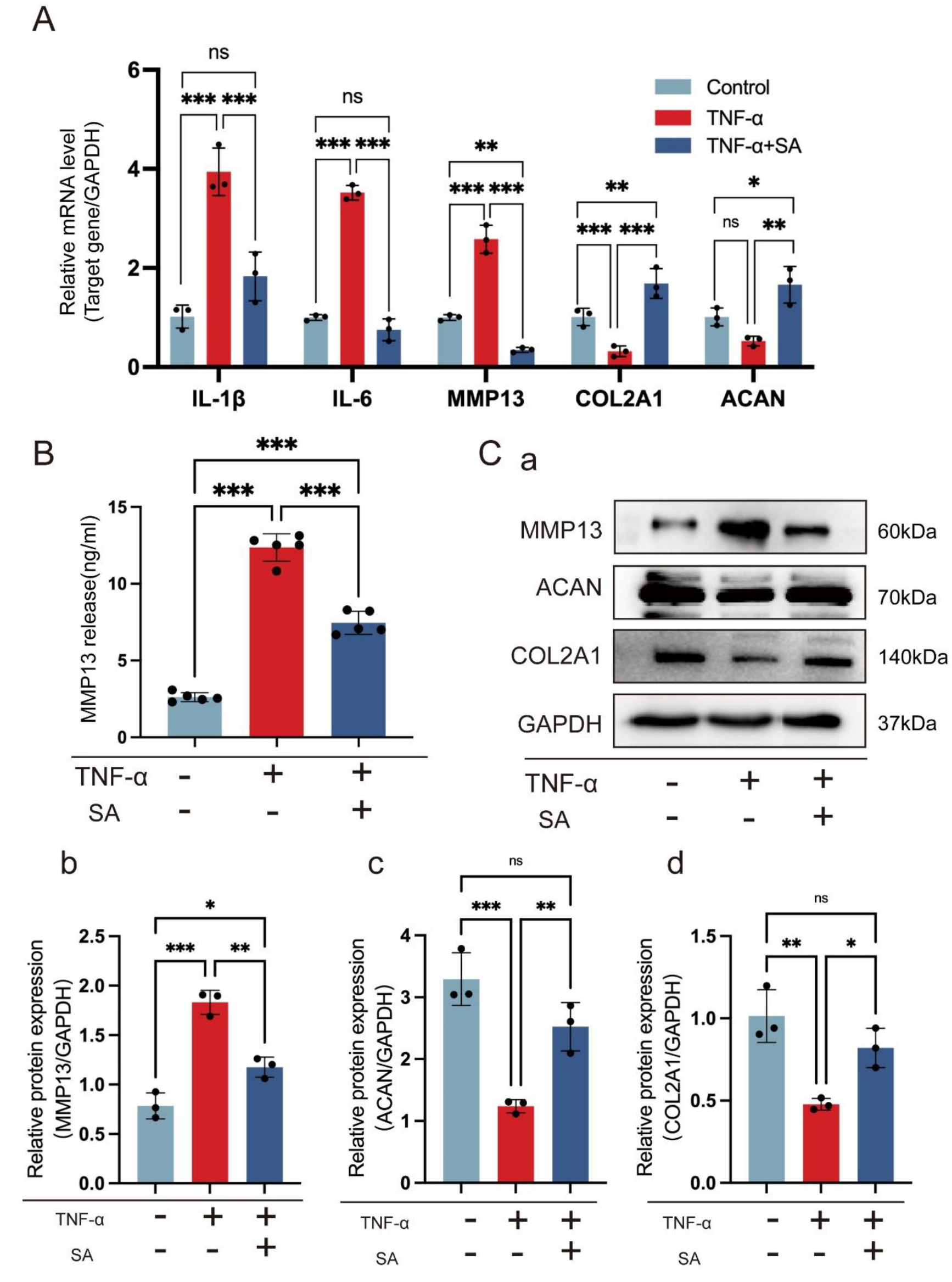
SA inhibits TNF-α induced chondrocytes inflammatory factors expression and extracellular matrix degeneration. Chondrocytes were stimulated with TNF-α and then treated or not treated with 10 μM SA for 48 hrs. (A) Relative mRNA expression levels of IL1β, IL-6, MMP13, COL2A1 and ACAN were detected by RT-qPCR (n=3, one-way ANOVA). (B) Expression of active MMP13 protein was reduced by the chondrocytes when stimulated with SA (n=3, one-way ANOVA). (C) WB analysis for detecting cartilage matrix markers MMP13, ACAN and COL2A1 in different groups (a). Quantitative analysis of MMP13(b), ACAN (c) and COL2A1(d) in protein level, GAPDH was used as reference protein (n=3, one-way ANOVA). The data are expressed as mean ±SD, *p<0.05, **p<0.01, ***p<0.001, and ns, not significant.

### 3.3 SA ameliorated cartilage degeneration by enhancing chondrocytes proliferation and inhibiting chondrocyte apoptosis

We next investigated the effects of SA on TNF-α mediated chondrocytes apoptosis and proliferation. We found TNF-α significantly inhibited antiapoptotic gene BCL-2 and promoting cell cycle genes CDK1 and MKi67 in chondrocytes. However, when chondrocytes were treated with 10 μM SA simultaneously, these inhibiting effects could be eliminated (Figure 3A). As for the phosphorylation of ERK, we found that TNF-α did not influence the total ERK (t-ERK) expression nor p-ERK statistically, however, SA could elevate the ratio of p-ERK and t-ERK compared with TNF-α treatment group (Figure 3B). Meanwhile, we determined the effects of SA on TNF-α mediated inhibiting of BCL2 and CDK1 expressions at the protein level, we found that SA eliminated these inhibiting effects mediated by TNF-α as well (Figure 3Ca). Quantitative analysis showed the same trend (Figure 3Cb-c). These results suggested that SA eliminated TNF-α mediated inhibition of cell cycle related markers expression.

**Figure 3:**
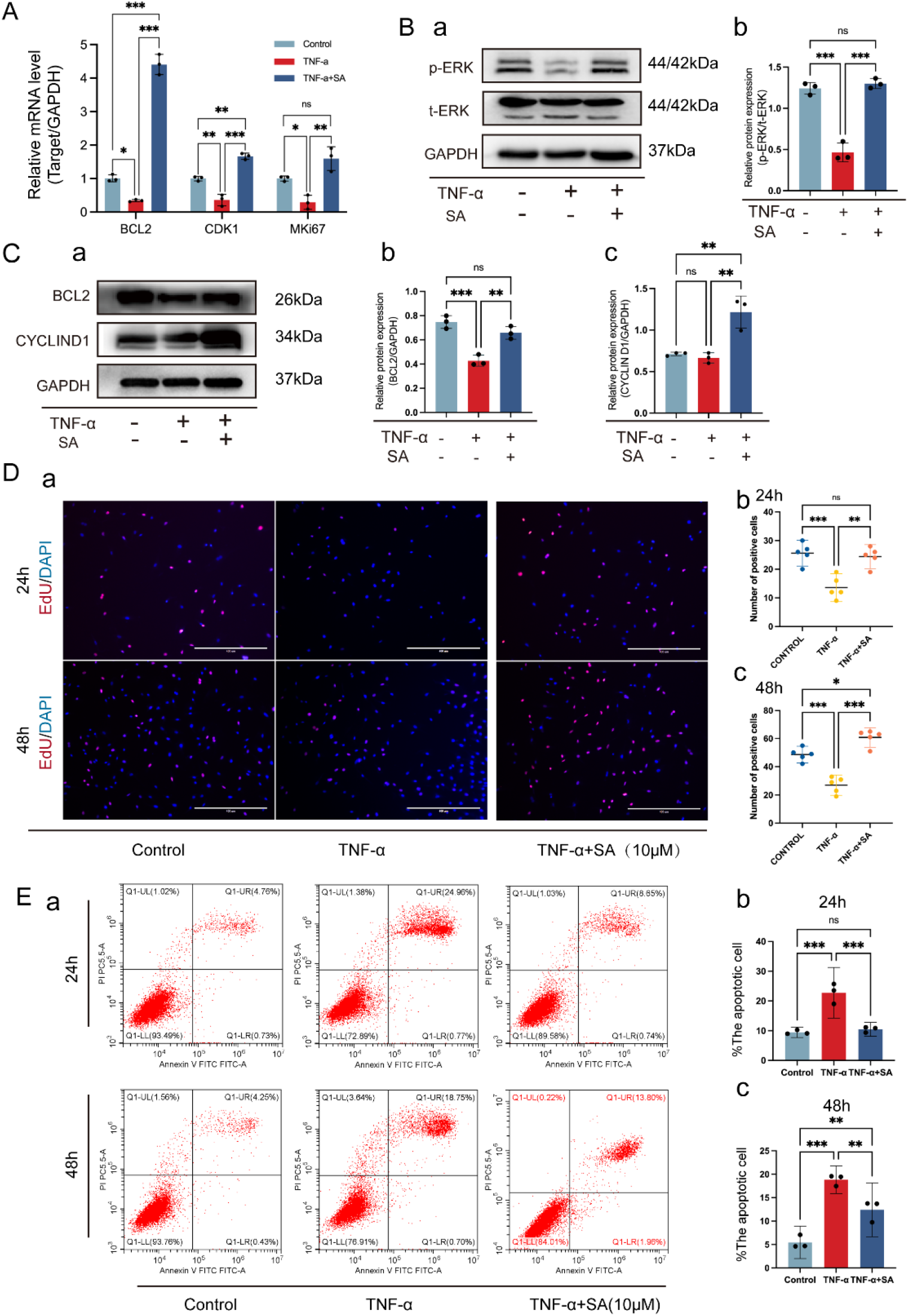
SA rescues TNF-α induced inhibition of chondrocytes proliferation and TNF-α induced promotion of chondrocyte apoptosis. (A) SA rescues TNF-α induced apoptosis markers expression at mRNA level. Chondrocytes were stimulated with TNF-α and then treated or not treated with 10 μM SA for 48 h, relative mRNA expression levels of BCL2, CKD1 and MKi67 were detected (n=3, one-way ANOVA). (B) Western blot analysis for detecting t-ERK and p-ERK in different group with the treatment of SA for 48 h(a). Quantitative analysis of ratio of p-ERK/t-ERK, GAPDH was used as reference protein (b) (n=3, one-way ANOVA). (C) WB for detecting cycle proliferation markers BCL2 and CYCLIN D1 in different group 48 hours after treatment at protein level. Quantitative analysis of BCL2 (b) and CyclinD1 (c) at protein level, GAPDH was used as reference protein (n=3, one-way ANOVA). (D) Fluorescent staining of EdU assay for detecting DNA synthesis indicating cell proliferation with 24 h and 48 h treatment (a), scale bar 400 μm. Percentage of EdU (red) positive staining statistical analysis at 24hrs (b) and 48hrs (c), (n = 5, random fields, one-way ANOVA). (E) Flow cytometry for cell apoptosis analysis of chondrocytes in each treatment group at 24hrs and 48hrs (a). Percentage of apoptotic cell in each treatment group at 24hrs (b) and 48hrs (c) (n=3, one-way ANOVA). The data are expressed as mean ±SD, *p<0.05, **p<0.01, ***p<0.001, and ns, not significant.

Furthermore, cell proliferation by EdU staining showed SA could rescue TNF-α induced inhibition of DNA duplication in dose-dependent manner at both 24h and 48h (Figure 3Da). Quantitative analysis showed the same trend (Figure 3Db). Cell apoptosis and quantitative analysis also confirmed that SA eliminated TNF-α induced promotion of cell apoptosis at both 24h and 48h (Figure 3E).

### 3.4 DEGs identification and pathway enrichment analysis

All sequencing data are available through the NCBI Sequence Read Archive under the accession number PRJNA820304.To figure out the potential mechanisms, RNA-seq was performed in chondrocytes cultured with or without SA under TNF-α stimulation. As exhibited in Figure 4A-B, for differentially expressed genes (DEGs), we found that 94 DEGs were up-regulated, and 96 DEGs were downregulated. Then DEGs were subjected to GO and KEGG signaling pathway enrichment analysis (Figure 4C-D). It comes to our attention that DEGs were highly enriched in biological process (BP) through GO analysis which including response to ER stress and cellular response to oxygen levels etc. What was more, KEGG signaling pathway enrichment analysis showed that protein processing in endoplasmic reticulum (Figure 4C-D marked with blue) was one of the most highly enriched signaling pathways.

**Figure 4:**
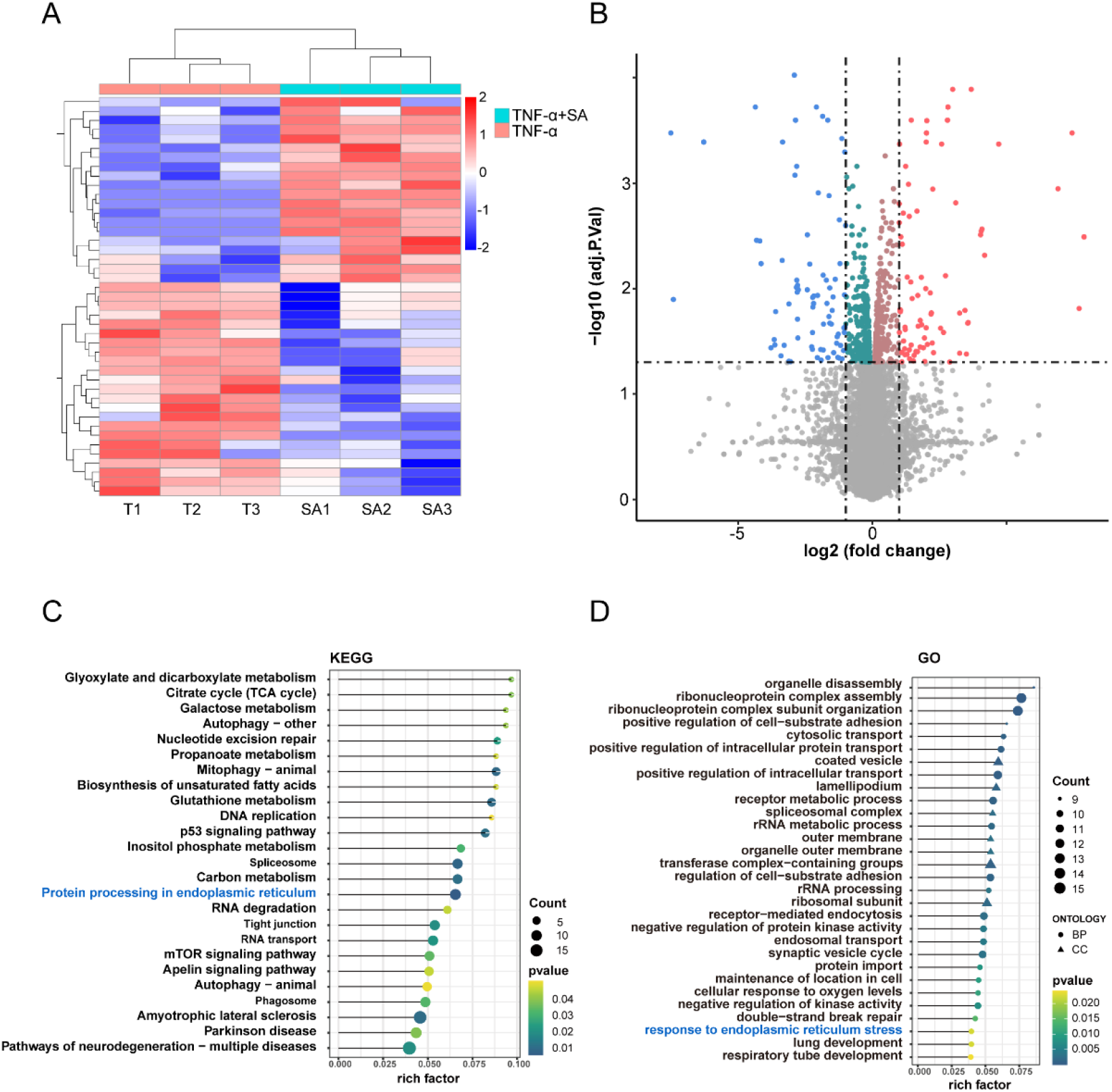
Bioinformatics analysis and popential mechanisms prediction. (A) Heatmap for global gene expression with group clusters (n=3). (B) Volcano map of DEGs in SA group vs control group (up-regulation: 75 genes and down-regulation: 32 genes), FC (fold change) ≥ 2 was accepted as positive DEGs. (C) Pathway enrichment bubble map based on the KEGG enrichment analysis, rich factor indicates a higher degree of enrichment, the larger p value (−log10) indicates a higher statistical significance, the larger bubble indicates a higher degree of enrichment. (D) GO enrichment of those DEGs, rich factor indicates a higher degree of enrichment, the larger p value (−log10) indicates a higher statistical significance, the larger bubble indicates a higher degree of enrichment, BP: biological process, CC: cellular component.

### 3.5 SA inhibiting IRE1α-IκBα-p65 signaling mediated ER stress by directly binding on IRE1α

As the ER transmembrane sensor, IRE1α is the key factor for ER stress. Therefore, we analyzed the potential combination of SA and IRE1α. Three dimensional (3D) structures of SA and IRE1α were subjected to molecular docking. The score of SA docking with IRE1α was −8.567 kcal/mol, and the theoretical binding mode was shown in Figure 5A. As shown, the compound and the amino acid residue TYR628 formed a hydrogen bond with a bond length of 3.3 Å, the amino acid residue CYS645 formed a hydrogen bond with a bond length of 3.2 Å, and the amino acid residue SER724 (phosphorylation site) formed a hydrogen bond with a bond length of 3.6 Å. These interactions guaranteed the stability of the combination of IRE1α and SA. Further, by DARTS assay we found that SA protected IRE1α from does dependent protease digestion compared with the control group (Figure 5B), which indicates the direct combination of SA and IRE1α protein. At the same time, ER-Tracker Red staining showed that TNF-α induced ER stress damage was rescued by SA treatment (Figure 5C). These results indicated that SA could binds on IRE1α and rescue TNF-α induced ER stress damage.

**Figure 5:**
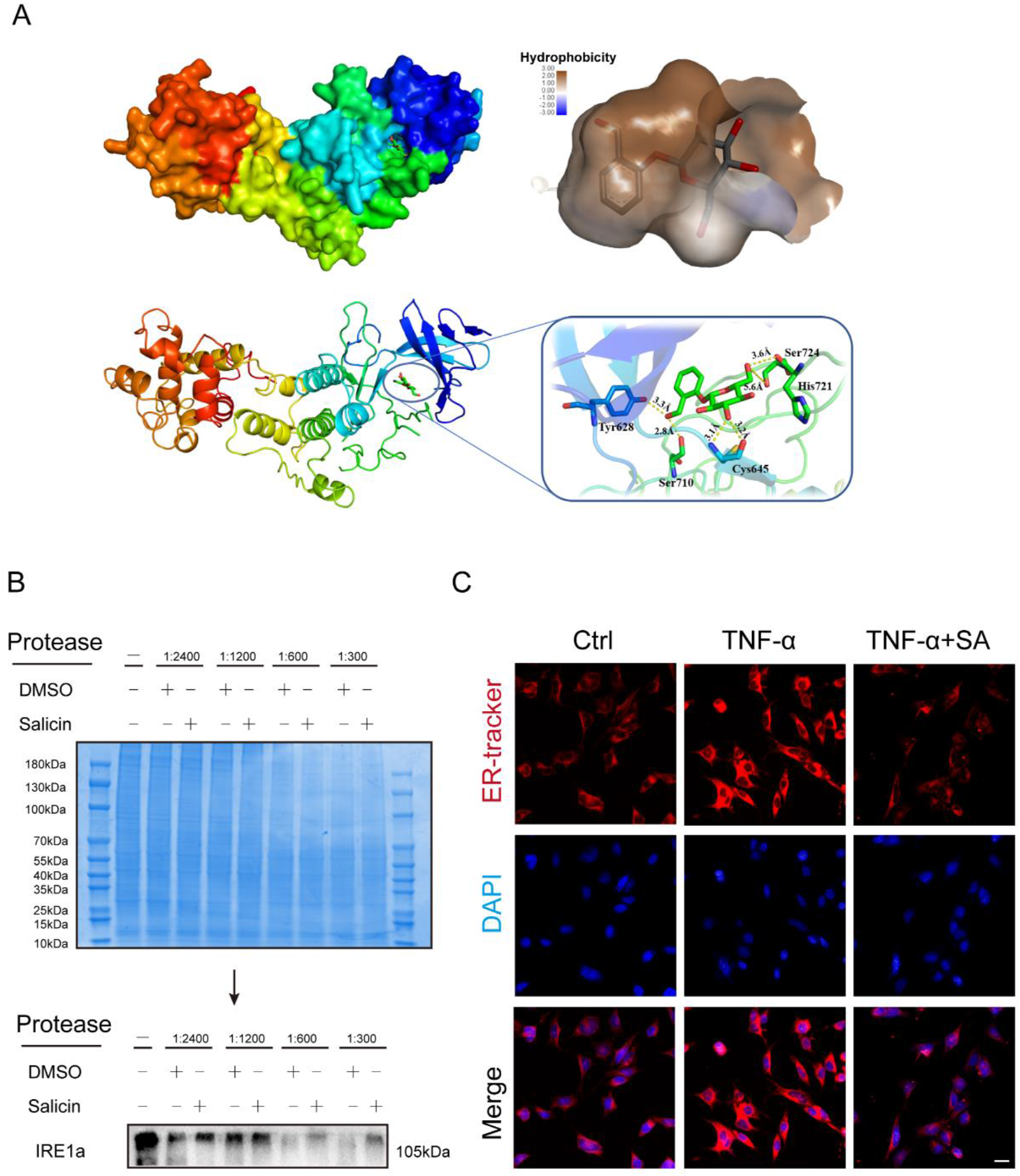
SA directly binded on IRE1α and inhibits IRE1α mediated ER stress. (A) Molecular docking for estimating combination site of IRE1α and SA. Three-dimensional (3D) structure of IRE1α and SA were subjected for molecular docking, combination site was obtained according to the software scoring system. Upper left panel: SA was located in the cavity formed by IRE1α 3D structure, upper right panel: magnification of the cavity; lower left panel: combination site of SA and IRE1α, lower left panel: magnification shown the hydrogen bonds (yellow dot line) formed between SA and IRE1α. (B) DARTS assay for confirmation of IRE1α and SA combination, protease was used to digest IRE1α protein, with the addition of SA, the digestion of IRE1α was significantly blocked at each concentration of protease compared with DMSO group.(C) Illustrative live-cell fluorescence microscopy images of ER damge labeled with ER-Tracker Red, chondrocytes exposed to TNF-α were treated with or without SA, ER damage was stained with red (upper panel), cell nucleus wree stained with DAPI (middle panel) and merged images were shown (lower panel).(scale bar: 50 μm).

Next, we investigated the regulation of SA on IRE1α mediated ER stress. Firstly, we detected the expression level of ER stress marker GRP78, and found SA inhibited TNF-α initiated GRP78 expression by WB (Figure 6Aa) and quantitative analysis (Figure 6Ab). Total IRE1α and p-IRE1α were also detected by WB, compared with control group, TNF-α did not influence total IRE1α expression, but downregulated phosphorylated IRE1α expression, which could be rescued by the treatment of SA (Figure 6Ac). Then we detected IRE1α downstream gene NF-kappa-B inhibitor alpha (IκBα) and p65 expression at protein level. The results showed that TNF-α induced high expressions of p-IκBα and p65 could be diminished by SA treatment (Figure 6Ad-e).

**Figure 6:**
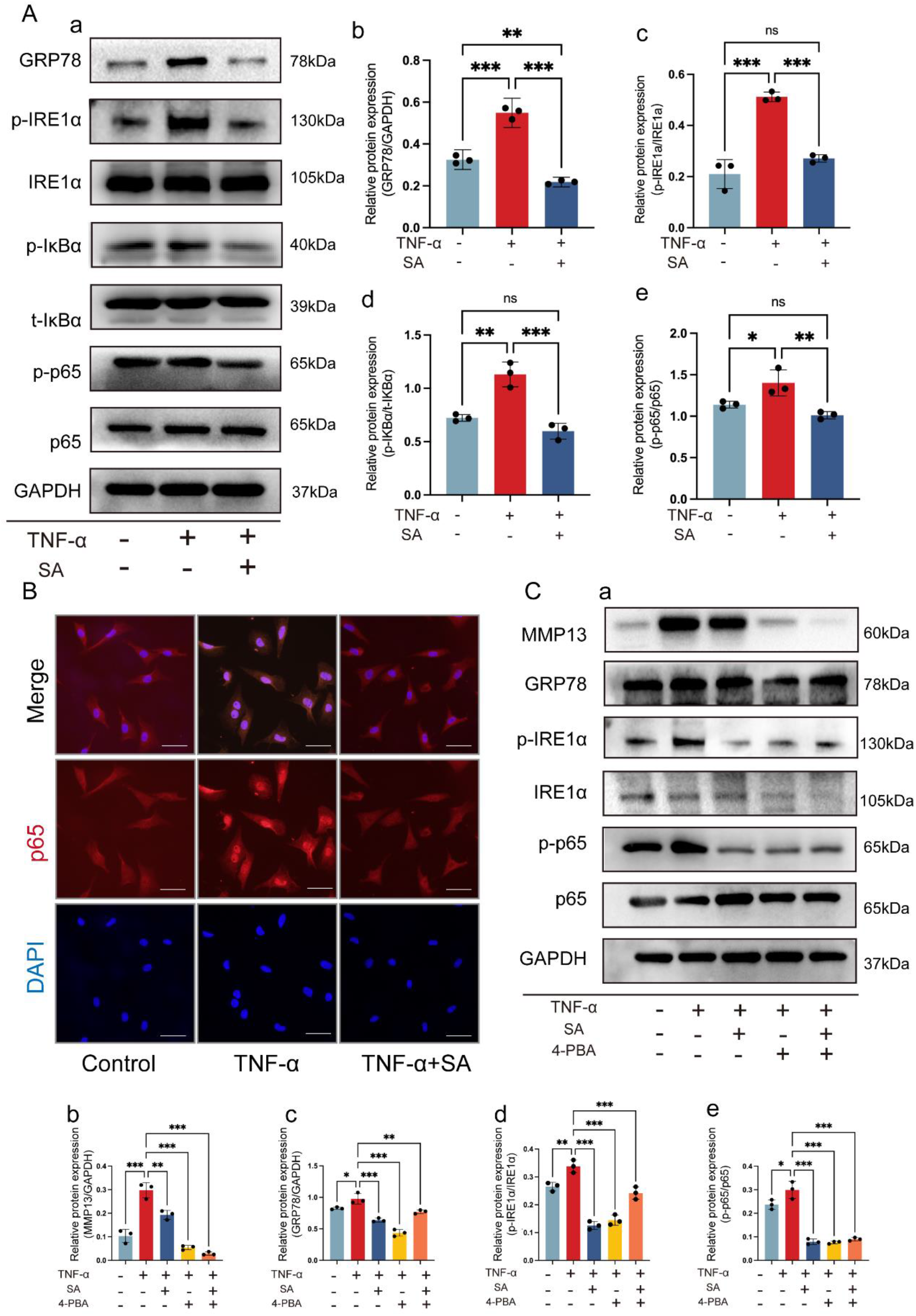
SA regulates IRE1α mediated ER stress by IRE1α-IκBα-p65 signaling. (A) WB analysis for detecting IRE1α and downstream genes expression at protein level. ER stress associated protein GRP78, pIRE1α, IRE1α, p-IκBα, t-IκBα, p-p65 and p65 were detected in each group (a). Quantitative analysis of GRP78 (b), ratio of p-IRE1α/t-IRE1α (c), ratio of p-IκBα/t-IκBα (d) and ratio of p-p65/t-p-p65 (e) at protein level, GAPDH was used as reference protein (n=3, one-way ANOVA). (B) P65 nucleus translocation in each treatment group. IF was used to detect the p-65 in nucleus, DAPI was used to staining cell nucleus, scale bar: 50 μm. (C) ER stress inhibitor 4-PBA blocked TNF-α mediated ER stress. Chondrocytes were stimulated with TNF-α and then treated with 10 μM SA or 15 μM 4-PBA for 48 h, MMP13, GRP78, IRE1α, p-IRE1α, p-p65 and p65 were detected by WB analysis (a). Quantitative analysis of MMP13 (b), GRP78 (c), ratio of p-IRE1α/t-IRE1α (d), and ratio of p-p65/t-p65 (e) at protein level, GAPDH was used as reference protein (n=3, one-way ANOVA). The data are expressed as mean ± SD, *p<0.05, **p<0.01, ***p <0.001, and ns, not significant.

Subsequently, we explored whether SA influenced TNF-α induced localization distribution of p65 protein in chondrocytes by IF. As expected, TNF-α induced p65 translocated from the cytoplasm into the nucleus were eliminated by SA treatment (Figure 6B). These findings indicated that SA inhibited IRE1α-IκBα-p65 signaling mediated ER stress by occupying the phosphorylation site of IRE1α.

Lastly, ER stress inhibitor 4-PBA was utilized for confirmation the function of SA. We found that 4-PBA dramatically inhibited TNF-α induced ER stress marker GRP78, p-IRE1α, p-p65 and cartilage matrix degeneration marker MMP13 expressions at protein level, and SA exhibited the similar function (Figure 6C). Taken together, these results suggested that SA directly could bind on IRE1α and inhibited IRE1α phosphorylation, which further blocked IRE1α-IκBα-p65 signaling mediated ER stress.

### 3.5 SA intra-articular injection ameliorates ACLT induced OA progression by inhibiting IRE1α mediated ER stress

Rat ACLT induced OA model was constructed for further research. A diagram summarized the progress of in *vivo* study (Figure 7A), SA loaded PLGA scaffolds were intraarticular injected for controlled release SA as reported previously^16,17^. As shown in Figure 7Ba, no obvious cartilage degeneration was found in the sham group. In ACLT and Vehicle groups, cartilage cell number, cartilage matrix and cartilage thickness decreased obviously compared with the sham group, which would be diminished in SA group. Knee joint OARSI scoring (Figure 7Bb) and quantitative analysis of articular cartilage area (Figure 7Bc) showed that SA treatment could reverse ACLT induced cartilage degeneration.

**Figure 7:**
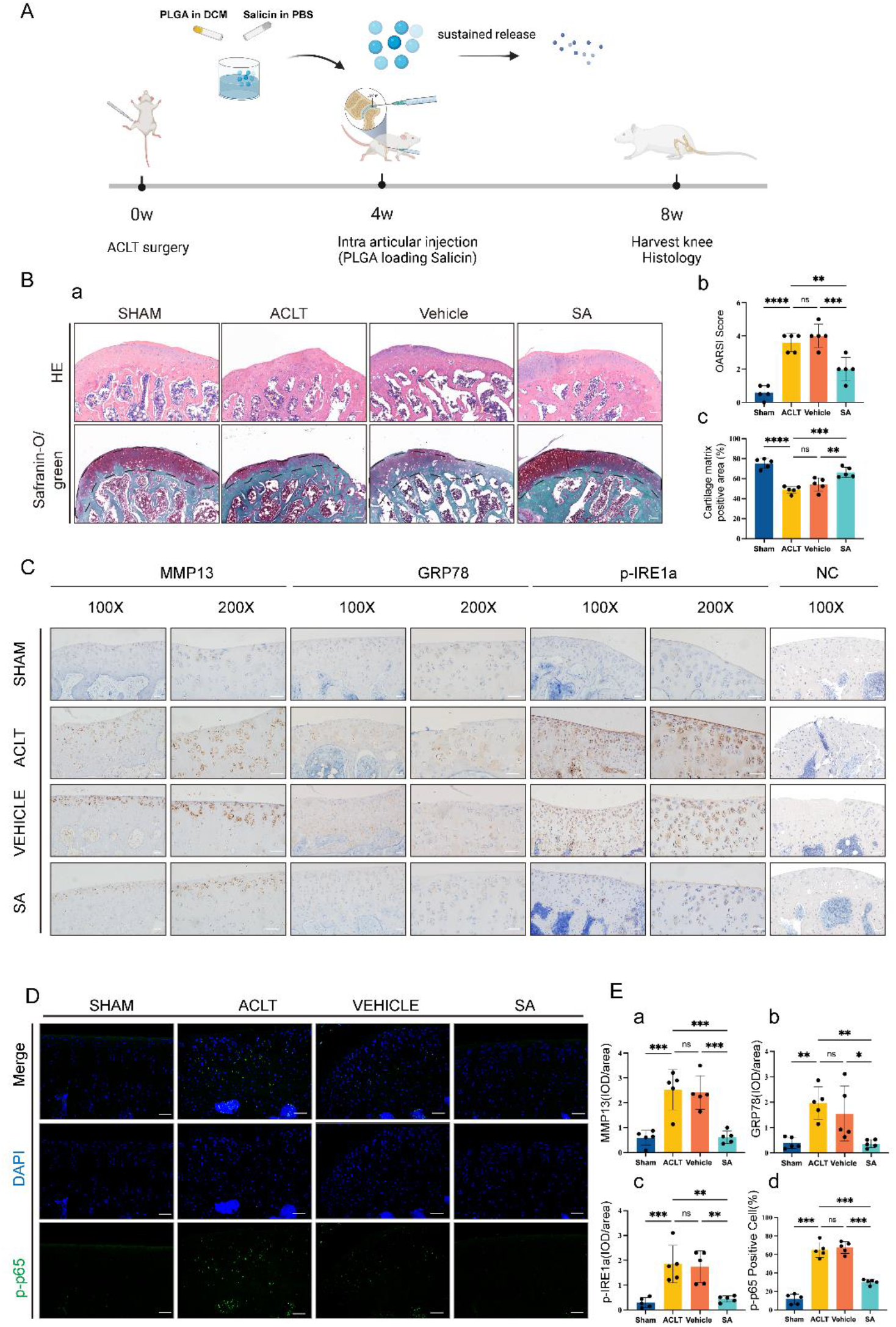
SA intra-articular injection ameliorates ACLT induced OA progression by inhibiting IRE1α mediated ER stress. (A) Diagram summarized the animal experiments. ACLT was used for the construction of knee OA model. Four weeks later, intra-articular injection of SA-loaded PLGA was done, and intra-articular injection of PLGA only was used as control. Four weeks after treatment, rats in each group were sacrificed and knee joints were subjected to histological analysis respectively. (B) H&E and Safranin-O/green staining for each treatment group. H&E (upper panel) and Safranin-O/green staining (lower panel) showed that ACLT resulted in the thicker of cartilage and degeneration of cartilage matrix compared with sham group, and intra-articular injection of SA-loaded PLGA ameliorated this progression compared with vehicle (PLGA only) group or sham group. OARSI scoring (b) and cartilage matrix area quantitative analysis (c) showed that, although OARSI score in ACLT, vehicle and SA groups were statistically higher than sham group, OARSI score in SA group significantly lower than ACLT and vehicle groups (n=5, one-way ANOVA). (C) IHC for detecting cartilage matrix degeneration MMP13, ER stress marker GRP78 and p-IRE1α, NC, negative control, scale bar 50 μm. (D) P-p65 nucleus translocation in each treatment group, scale bar 50 μm. (E) Quantitative analysis showed that the expression of MM13 (a), GRP78 (b) and p-IRE1α (c) in ACLT group or vehicle group were significantly more than that in sham group, and the expression of MMP13, GRP78 and p-IRE1α in SA group were significantly less than those in ACLT group and vehicle group (n=5, one-way ANOVA). Quantitative analysis showed that P-p65 nucleus translocation in ACLT group or vehicle group was significantly more than that in sham group, and p-p65 nucleus translocation in SA group was obviously less than that in ACLT group and vehicle group (d) (n=5, random fields, one-way ANOVA). ACLT group indicates the group with PBS injection; Vehicle group indicates only inject PLGA vehicle; SA group indicates injection of SA loaded PLGA. Dash lines indicate cartilage surface. The data are expressed as mean ±SD, *p<0.05, **p<0.01, ***p<0.001, and ns, not significant.

To further prove the mechanisms of SA inhibiting IRE1α mediated ER stress, IHC and IF were carried out. We first detected that MMP13 was highly expressed in the articular cartilage matrix area in ACLT and vehicle groups compared with the sham group, and SA treated group decreased the expression of MMP13 in the cartilage matrix area. Then, we determined the ER stress marker GRP78 expression, we found that OA induced a high expression of GRP78 in cartilage cell could be reversed by SA treatment. Thirdly, we found OA induced high phosphorylation level of IRE1α was alleviated by SA treatment (Figure 7C). Finally, we identified that OA induced p-p65 nuclear translocation was inhibited by SA treatment (Figure 7D). Quantitative analysis of IHC and IF were shown in Figure 7Ea-d. These results suggested SA ameliorates ACLT induced OA progression by inhibiting IRE1α mediated ER stress in *vivo*.

In summarize, our in *vitro* and in *vivo* tests showed that SA bound on IRE1α and blocked IRE1α phosphorylation, then inhibits IRE1α mediated ER stress by IRE1α-IκBα-p65 signaling (Figure 8).

**Figure 8:**
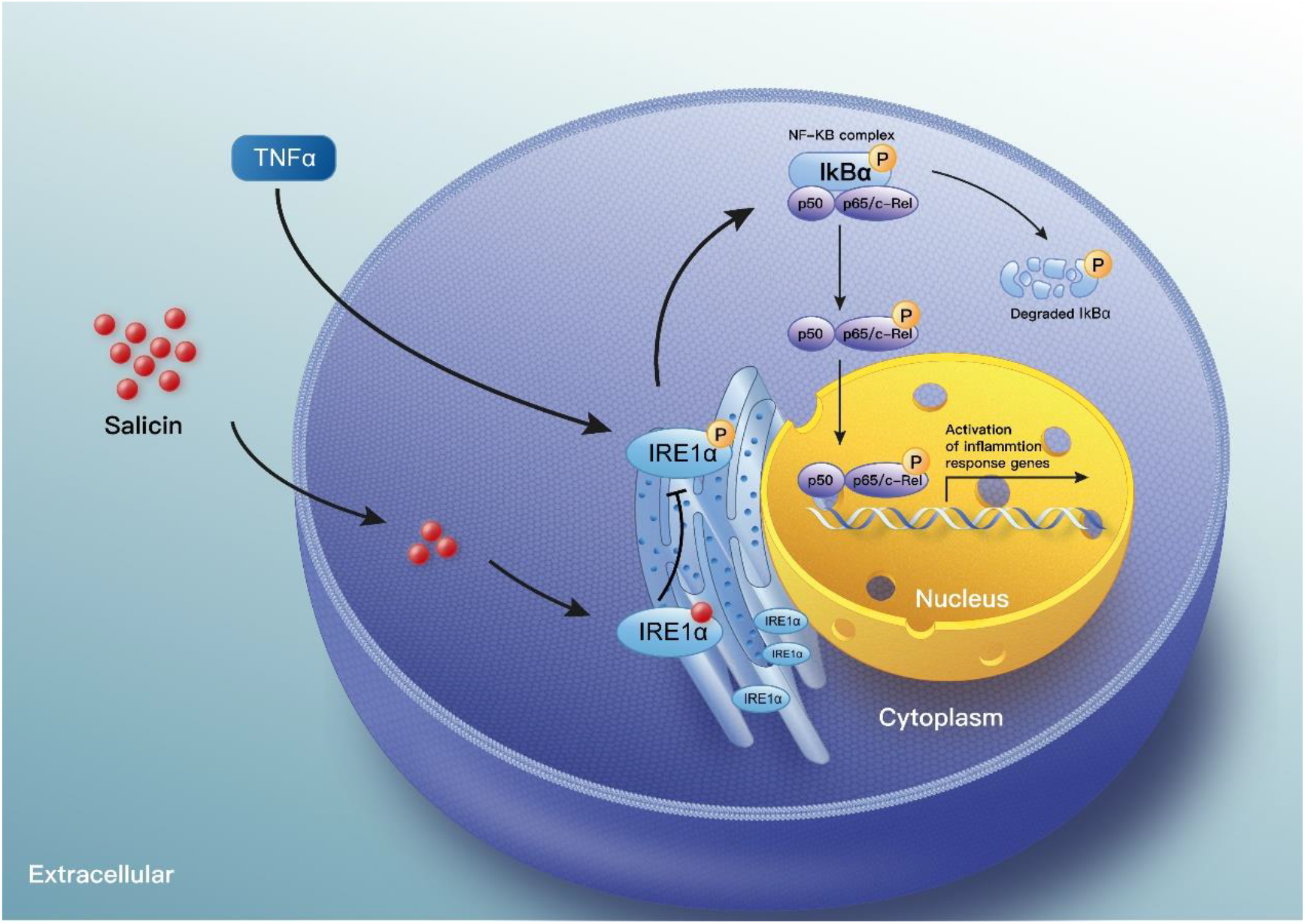
A proposed model of action. SA bound on IRE1α and blocked IRE1α phosphorylation, then inhibits IRE1α mediated ER stress by IRE1α-IκBα-p65 signaling.

## 4. Discussion

With the population aged, the morbidity of OA is increasing year by year^24^. The discovery and development of DMOADs have been identified as potential methods for ameliorating the effect of increasing OA prevalence^2,25^. In the present study, we identified SA not only inhibits TNF-α induced chondrocytes inflammatory factor expression and extracellular matrix degeneration but also enhances chondrocytes proliferation and inhibits chondrocytes apoptosis. In mechanism, we clarified SA bound with IRE1α directly and blocked IRE1α mediated downstream ER stress.

As the main chemically standardized of willow bark extract, SA also belongs to non-steroidal anti-inflammatory drugs (NSAIDs), which may cause gastrointestinal, renal, and cardiovascular toxicity^6,25^. Therefore, topical use should be considered rather than oral administration. As a natural small molecular drug, SA can cross the cell membrane and bound with intracellular targets, thus relieve serious side effects caused by oral administration. By directly adding SA to chondrocytes medium and intra-articular injection of SA loaded PLGA, we demonstrated the anti-inflammatory, promotes proliferation and anti-apoptosis effect of SA in the progression of OA. These results indicate the efficiency of intra-articular use of SA.

Recently, except the anti-inflammatory, antipyretic, antirheumatic, and antiseptic properties^10^, Gao et al^26^ found that SA prevents collagen type II degradation by inhibiting the activation of NF-κB proinflammatory pathway. However, the underlying mechanisms in detail are still unclear. In current study, we obtained similar results by detecting cartilage matrix degeneration and inflammatory markers. Furthermore, DGEs found by RNA sequencing and KEGG enrichment analysis showed that ER stress was highly involved in SA mediated protective effects of cartilage degeneration. Next, molecule docking and DARTS analysis showed SA could occupy the phosphorylation sites of IRE1α and block IRE1α phosphorylation mediated p65 nucleus translocation and downstream gene expression. Thus, the current study applied a new field of version for understanding SA mediated protective effects of cartilage degeneration.

The ER is a multifunctional organelle, where protein folding occurs prior to transport to extracellular surface or to different intracellular sites^27-30^. Three ER transmembrane proteins mediated UPR, including IRE1, pancreatic endoplasmic reticulum kinase (PERK), and activating transcription factor 6 (ATF6) ^28,29^. Among them, IRE1 is the most conserved gene from yeast to human, and it has two subtypes: IRE1α and IRE1β. IRE1α is commonly expressed in most cells and tissues, while IRE1β is limited to gastrointestinal epithelial cells^31^. Worth mention, IRE1α plays key role in chondrocytes proliferation, ECM production etc.^32^, which indicates the regulatory function of IRE1α mediated ER stress in chondrocytes proliferation and degeneration. Jacqueline et al^33^ found two top risk factors for OA, age and obesity, were highly associated with ER stress, and resveratrol could mitigate early joint degeneration by inhibiting ER stress. Benedetta et al^34^ found that chronic ER stress decreased chondrocyte proliferation. Kung et al^35,36^ reported that hypertrophic chondrocytes hold limited potential to cope with increased ER stress, they also found increased ER stress is sufficient to reduce chondrocyte proliferation. Rajpar et al’s research identified that ER stress play a direct role in cartilage pathology^37^. Meanwhile, it is widely accepted that ER stress is directly associated with chondrocytes apoptosis and death^38-40^. What’s more, recent studies confirmed OA inflammation is also related to ER stress^41-45^. Be similar with these obtained conclusions, we found by directly bound with IRE1α, SA inhibited IRE1α mediated ER stress and subsequently promotes chondrocytes proliferation, decreases inflammatory factors expression and inhibits chondrocytes apoptosis.

Recently, ER stress regulators were identified as new drug targets for several diseases^27-29,46^. During the progression of OA, chondrocytes are responsible for the biogenesis and maintenance of cartilage ECM. Several cellular stresses including hypoxia, oxidative stress, nutrient deprivation, aging or injury etc. could cause excessive unfolding or misfolded proteins on ER and trigger ER stress^33,40^. Huang et al^32^ reported that IRE1α regulates chondrocytes apoptosis by activating NF-κB signaling. Ye et al^47^ reported phosphorylated IRE1α activating NF-κB signaling by releasing IκBα. Released NF-κB dimers translocated to the nucleus and binded κB sites in the promoters or enhancers of target genes^48^. We found in the progression of OA, ER stress was mediated by p-IRE1α, p-IκBα and activating p-p65 nucleus translocation. With SA treatment, this process was inhibited. Our molecule docking and DARTS analysis showed that SA can bind the phosphorylation site of IRE1α and block IRE1α phosphorylation, followed by the decreasing expression of p-κBα and p-p65 nucleus translocation. Therefore, as one of the key ER stress regulators, IRE1α is the potential drug targets for OA treatment.

As whole joint disease, OA especially late-stage OA characterized with dramatically synovium, cartilage and subchondral bone pathology^1,49,50^. However, early-stage OA was featured with cartilage pathology^51,52^, SA may be an effect drug for modifying early-stage OA. Further studies focus on the effect of SA on the other cell types such as synoviocytes, immune cells etc. will contribute to further clinical use of SA. Collectively, our results have demonstrated SA is directly bound on IRE1α and blocks IRE1α phosphorylation, then inhibits IκBα phosphorylation and p65 nucleus translocation, finally inhibits chondrocytes apoptosis, promotes chondrocytes proliferation and decreases inflammatory factors expression. Thus, SA is a potential drug for clinical modifying OA progression by intra-articular injection.

## Acknowledgments

We are grateful for the support of Key Laboratory of Biorheological Science and Technology, College of Bioengineering, Chongqing University. The authors would like to thank Dr. Tingting Peng from the Chongqing BI academy.

## Contributions

Conception and design: Wei Huang and Junyi Liao; Analysis and interpretation of the data: Zhenglin Zhu, Shengqiang Gao, Cheng Chen, Wei Xu, Pengcheng Xiao, Chengcheng Du, Bowen Chen, Yan Gao; Drafting of the article: Junyi Liao, Zhenglin Zhu; Critical revision of the article for important intellectual content: Junyi Liao and Wei Huang; Provision of study materials: Junyi Liao, Chunli Wang and Wei Huang; Statistical expertise: Zhenglin Zhu; Obtaining of funding: Junyi Liao, Zhenglin Zhu, and Wei Huang; Administrative, technical, or logistic support: Junyi Liao and Wei Huang; Collection and assembly of data: Zhenglin Zhu, Junyi Liao, Zhiyu Chen and Chunli Wang; Final approval of the article: all authors.

